# GIMLET: Identifying Biological Modulators in Context-Specific Gene Regulation Using Local Energy Statistics

**DOI:** 10.1101/349928

**Authors:** Teppei Shimamura, Yusuke Matsui, Taisuke Kajino, Satoshi Ito, Takashi Takahashi, Satoru Miyano

**Affiliations:** Division of Systems Biology, Nagoya University Graduate School of Medicine, 65 Tsurumai-cho, Showa-ku, Nagoya 466-8550, Japan; Laboratory of Intelligence Healthcare, Nagoya University Graduate School of Medicine, 1-1-20 Daiko-Minami, Higashi-ku, Nagoya 461-8673, Japan; Division of Molecular Carcinogenesis, Nagoya University Graduate School of Medicine, 65 Tsurumai-cho, Showa-ku, Nagoya 466-8550, Japan; Human Genome Center, Institute of Medical Science, The University of Tokyo, 4-6-1 Shirokane-dai, Minato-ku, Tokyo 108-8639, Japan

**Keywords:** Gene regulation, Modulator detection, Energy statistics, Distance correlation, Statistical test

## Abstract

The regulation of transcription factor activity dynamically changes across cellular conditions and disease subtypes. The identification of biological modulators contributing to context-specific gene regulation is one of the challenging tasks in systems biology, which is necessary to understand and control cellular responses across different genetic backgrounds and environmental conditions. Previous approaches for identifying biological modulators from gene expression data were restricted to the capturing of a particular type of a three-way dependency among a regulator, its target gene, and a modulator; these methods cannot describe the complex regulation structure, such as when multiple regulators, their target genes, and modulators are functionally related. Here, we propose a statistical method for identifying biological modulators by capturing multivariate local dependencies, based on energy statistics, which is a class of statistics based on distances. Subsequently, our method assigns a measure of statistical significance to each candidate modulator through a permutation test. We compared our approach with that of a leading competitor for identifying modulators, and illustrated its performance through both simulations and real data analysis. Our method, entitled genome-wide identification of modulators using local energy statistical test (GIMLET), is implemented with R (≥ 3.2.2) and is available from github (https://github.com/tshimam/GIMLET).

## 1 Introduction

The regulation of gene expression is a process in which the expression of a particular gene can be either activated or repressed. Transcription factors (TFs) contribute greatly to the process of gene regulation by binding to a specific DNA sequence in the promoter regions of their target genes and controlling their transcription. The responsiveness of a target gene expression to a TF typically varies owing to genetic variation or a change in the cellular environment. This modulation in gene-specific responsiveness is often caused by a specific factor, called a modulator, at different levels, including the transcriptional, post-transcriptional and post-translational levels.

In the last decade, large international consortia, such as The Cancer Genome Atlas (TCGA) [1] and the International Cancer Genome Consortium (ICGC) [2], have generated large-scale gene expression profiles of different tumor types and catalogued their genetic alterations (recurrent mutations and copy number variations). Genome-wide association studies (GWAS) have also identified tens of thousands of human disease-associated variants and millions of single nucleotide polymorphisms [3]. However, it remains unknown if and in what way many genetic alterations and variants interact with physical and functional interactions within cellular networks.

The identification of genetic alterations and variations that function as biological modulators and contribute to gene expression control is one of the challenging tasks in systems biology. Recently, sophisticated algorithms have been developed for this task, which has successful applications in many areas [4,5,6,7,8,9]. For example, MINDy [4] formulated the problem of identifying modulators as a problem of testing whether the expressions of a univariate TF and its target gene, denoted by *X* and *Y*, are independent each other, conditioned on the expression levels of an univariate modulator denoted by *Z* in the framework of conditional mutual information. GEM [5] used a linear regression model with the effects of interaction between *X* and *Z* to describe the relationships between *X* and *Y* modulated by *Z*. MIMOSA [6] considered a mixture model of X and *Y* from two different fractions based on *Z*. Note that these methods were designed to capture a particular type of three-way dependency, where *X*, *Y*, and *Z* are univariate random variables. Therefore, they cannot capture multivariate dependencies where sets of random variables are associated with each other. Currently, no systematic mathematical framework exists for identifying biological modulators of complex gene regulation, such as combinatorial regulation, where multiple TFs and modulators are functionally related.

In this study, we present a novel method, genome-wide identification of modulators using local energy statistical test (GIMLET), to overcome the challenges outlined above. GIMLET includes the following contributions.

1. GIMLET is mainly based on dependence coefficients from energy statistics for modeling the relationships between genes. These types of coefficients are a measure of the statistical dependence between two random variables or two random vectors of arbitrary, not necessarily equal dimension. This enables the correlation of the expression of sets of any size for TFs, their target genes, and modulators.
2. We provide a new dependence coefficient, called local distance correlation, to compare the difference in distance correlation at low and high values of given modulators, allowing the identification of all types of local dependencies, such as nonmonotone and nonlinear relationships, between TFs and their target genes at the fixed point of modulators.
3. We develop a permutation-based approach to evaluate whether local distance correlation varies with modulators, which enables the discovery of modulators related to complex regulatory relationships, including synergistic and cooperative regulation, from a statistical point of view.

We describe our proposed framework and algorithm in Section 2. We present the efficiency of GIMLET using synthetic and real data in Sections 3 and 4.

## 2 GIMLET methodology

### 2.1 Notations and preliminaries

For a *p*-dimensional random vector ***a***, |***a***| represents its Euclidean norm. A collection of *n* independent and identically distributed (i.i.d.) observations for ***a*** is denoted as {***a****_k_*; *k* = 1,…,*n*} where ***a****_k_* = (*a*_1_,…,*a_p_*)′ represents the *k*-th sample.

The distance correlation [10] was introduced as a measurement of dependence between two random vectors *X* ∈ **R***^p^* and *Y* ∈ **R***^q^*. It is based on the concept of distance covariance between *X* and *Y*, denoted by 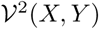, which measures the distance between the joint characteristic function of (*X*, *Y*) and the product of the marginal characteristic functions as follows:

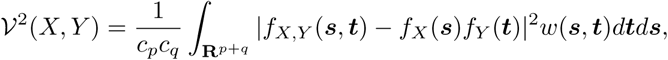

where *f_X,Y_* (***s***, ***t***), *f_X_* (***s***), and *f_Y_* (***t***) are the characteristic functions of (*X*,*Y*), *X*, and *Y*, respectively, and the weight function *w*(***s***, ***t***) = (*c_p_c_q_*|***s***|^1+^*^p^*|***t***|^1+^*^q^*)^−1^ with constants *c_l_* = *π*^(1+^*^l^*^)^*^/^*^2^*/Γ* ((1 + *l*)*/*2) for *l* ∈ **N** is chosen to produce a scale-free and rotation-invariant measure that does not go to zero for dependent variables.

The distance correlation 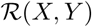 between *X* and *Y* is then defined as

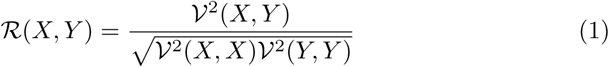

if 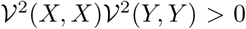 and equals 0 otherwise. The remarkable properties of the distance correlation introduced by the equation (1) include 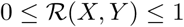 and 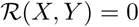 if and only if *X* and *Y* are independent.

If we observe a collection {(***x****_k_,* ***y****_k_*); *k* = 1,…,*n*} of *n* i.i.d. observations from the joint distribution of random vectors 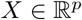 and 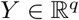, the empirical distance covariance between *X* and *Y*, denoted by 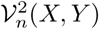, is then given by

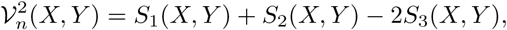

where

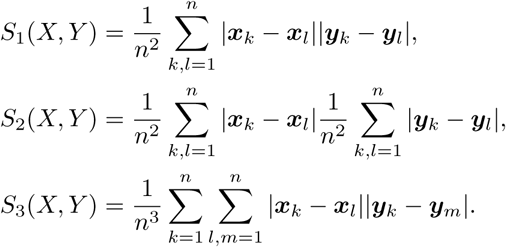

The empirical distance correlation 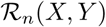 is then

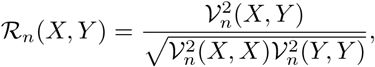

and satisfies 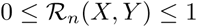.

### 2.2 Local distance correlation

We introduce a local estimator of the distance correlation evaluated at another random vector 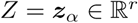 as a local measurement of the dependence between *X* and *Y* conditioning on *Z* = ***z****_α_* based on the observed data. We consider a collection {(***x****_k_,* ***y****_k_,* ***z****_k_*): *k* = 1,…,*n*} of *n* i.i.d. observations for random vectors *X*, *Y*, and *Z*. Let us denote *w_kα_* = *K_h_*(***z****_k_,* ***z****_α_*) satisfying 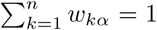 as the new weight function based on the distance between two sample vectors ***z****_k_* and ***z****_α_* where *K_h_* is a specified kernel function with a bandwidth *h*.

Based on the definition of the Nadaraya-Watson estimator [12,13] as a weighted averaging method, we define a local estimator of distance covariance conditioning on *Z* = ***z****_α_*, using the weighted Euclidean distance as

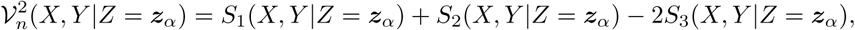

where

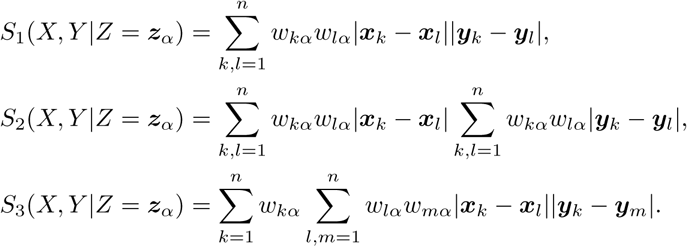

Each sample of the neighborhood in the *α*-th sample is weighted according to its weighted Euclidean distance from *Z* = ***z****_α_*. Points close to *Z* = ***z****_α_* have a large weight, and points far from *Z* = ***z****_α_* have a small weight. The kernel function *K_h_* used in all of our examples is the Gaussian kernel function *K_h_*(***z****_k_,* ***z****_α_*) = exp(−|***z****_k_* − ***z****_α_*|^2^/*h*) where *h* is a bandwidth parameter that controls the smoothness of the fit. For a specific point *Z* = ***z****_α_*, the nearest-neighbor bandwidth *h* is determined such that the local neighborhood contains the *q* = ⎿*nδ*⏌ closest samples to the *α*-th sample in the Euclidean distance of *Z*, where *δ* ∈ (0, 1) is a tuning parameter that indicates the proportion of neighbors. Therefore, each local estimator is inferred with *q* observations that fall within the sphere *B_δ_*(***z****_α_*), centered at the *α*-th sample. We use a varying width parameter *h* that reduces the problem of data sparsity by increasing the radius in the regions with fewer observations.

The empirical local estimator of the distance correlation, called local distance correlation, 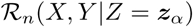 for given *Z* = ***z****_α_* is then defined by the equation

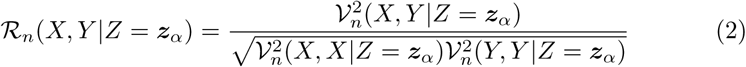

if both 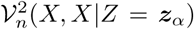 and 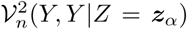 are strictly positive, and it is equal to zero otherwise.

### 2.3 Statistical hypothesis test for identifying modulators

In the statistical hypothesis testing for identifying modulators, it is of practical interest to assess whether the local dependence between *X* and *Y* varies with *Z*. This question can be formulated as:

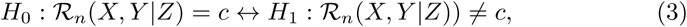

where 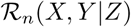 is a function of *Z* and *c* is a constant.

For calculating the *p*-values of the local dependence between *X* and *Y* for each *Z*, we apply a permutation-based approach similar to the one used by [4]. Under the assumption that 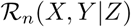 is a monotonic function of *Z*, we calculate the test statistic:

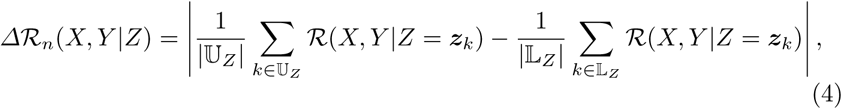

where 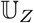 and 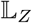 are the index sets of the upper and lower points of *Z*, respectively. To assess the statistical significance of 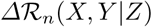, we generate a series of null hypotheses, and calculate the empirical *p*-value, using the following permutation procedures:

1. Permute the values of *Z* for all samples.
2. Re-calculate the test statistics using (4). Denote the null statistic of the *l*-th permutation by 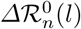.
3. Repeat steps 1-2 *B* times and calculate the empirical *p*-value for *Z*:

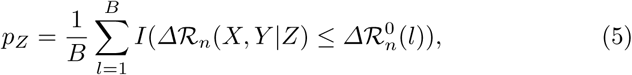

where the indicator function *I*(*A*) equals one when the condition *A* is true and it equals zero otherwise.

The statistical significance of *Z*, as expressed in (5), is the percentage of null statistics, equally or more extreme than the observed statistic for the given *Z*. Note that this empirical method directly couples both the minimal obtainable *p*-value and the resolution of the *p*-value to the number of permutations *B*. Therefore, it requires a very large number of permutations to calculate the *p*-values when we want to accurately estimate small *p*-values. In order to compute more accurate *p*-values, we use a semi-parametric approach based on a tail approximation [14,15]. The corrected empirical *p*-value 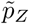, using the distribution tail approximation, is given by

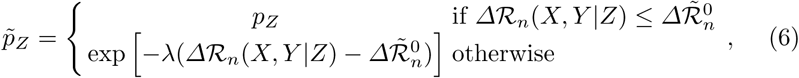

where *λ* is a scale parameter, and 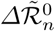 is a threshold that we set to the 99-th percentile of null distributions. The parameter *λ* is estimated by the null statistics satisfying the condition 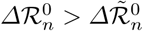.

## 3 Synthetic data results

We generated synthetic data and evaluated the performance of our method in order to gain insight into the statistical power and type I error rate control in identifying modulators, based on the hypothesis 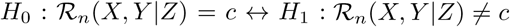.

A simulation study was conducted as follows. An i.i.d. sample of (*X, Y, Z*) was generated using the endogenous switching regression model under the following three settings:

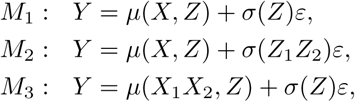

with

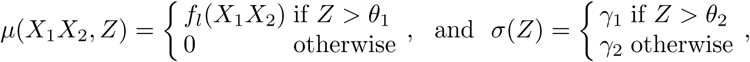

where *X*, *X*_1_, *X*_2_, *Z*, *Z*_1_, *Z*_2_ ~ *U*[0, 1], *ε* ~ *N*(0, 1), *µ* and *σ* are the conditional mean and variance of *Y* depending on *Z*, and *f_l_* is a function that determines a functional relationship between *X* and *Y*.

For a function *f_l_*(*X*), we considered the following eight functional relationships:

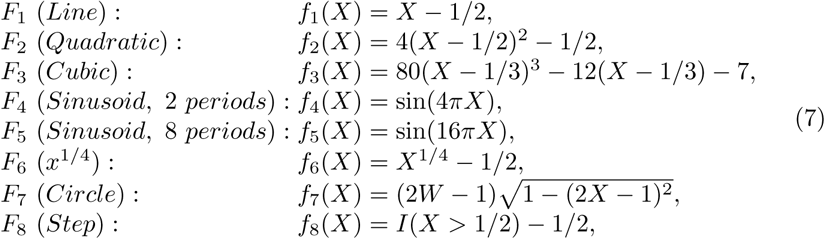

where *W* ~ *Bern*(0.5). These functions were originally used in [16] to assess the statistical power against independence.

We set *θ*_1_ and *θ*_2_ to be 0.25 and 0.75, respectively, and *γ*_1_ and *γ*_2_ as follows:

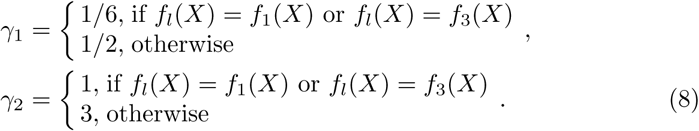

Scatter plots of the data obtained from these eight relationships are shown in Figure 1.

The first setting, *M*_1_, was designed to find modulators in the traditional framework for identifying modulators [4], where the expression value of a modulator 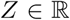 influences the dependence between the expression values of a TF 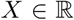 and its target gene 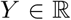. The second and third settings, *M*_2_ and *M*_3_, were aimed at finding the modulators in the new conceptual framework investigated in this study: *M*_2_ was intended for the combinatorial modulation, where the expressions of two modulators 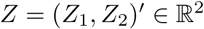 influence the dependency between a TF 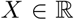 and its target gene 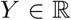. *M*_3_ was intended for combinatorial regulation, where the expression of a modulator 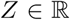 influences the dependency between two TFs 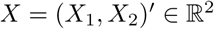 and their target gene 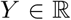, and both *X*_1_ and *X*_2_ are required for *Y*.

**Fig. 1.**
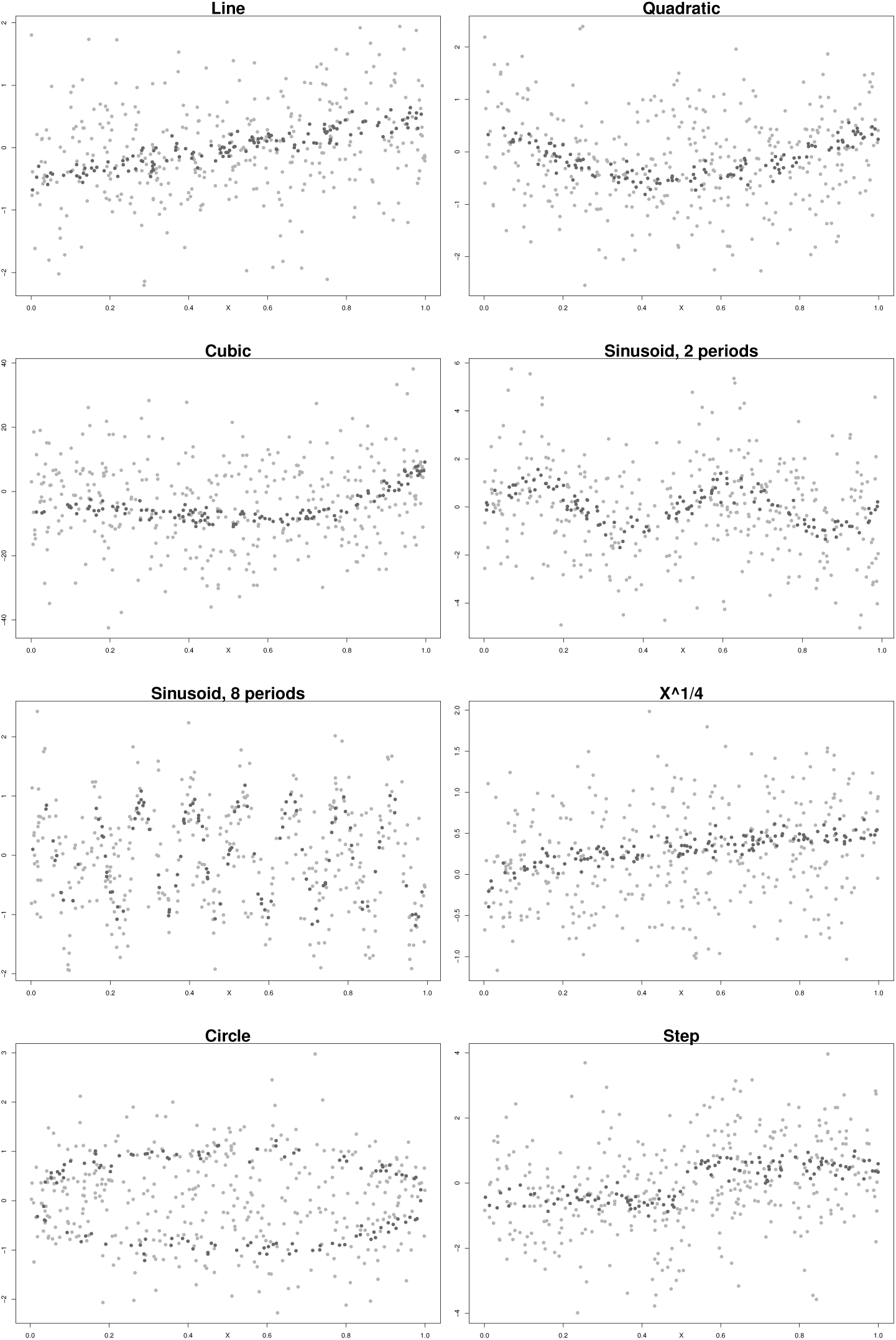
Sample plots of the eight simulated relationships. Dark gray dots indicate samples with *Z* > 0.75, whereas light gray dots indicate samples with *Z* ≤ 0.75.

The identification of modulators using our method (GIMLET) was assessed by comparing it with MINDy [4], one of the most widely used methods for this purpose. We note that MINDy cannot be directly applied to the identification of modulators under the settings *M*_2_ and *M*_3_, because MINDy is not designed for combinatorial modulation and regulation. In these simulations, all possible triplets were tested separately using MINDy, and the statistical significance was evaluated by using Fisher’s method, which is widely used to combine *p*-values. A hypothesis testing problem for identifying modulators with varying sample sizes (*n* = 100, 200, 500) was simulated with 1,000 datasets generated for each of the above three settings. All tests were performed at the significance level *α* = 0.05. The statistical power was estimated by the fraction of test statistics that were at least as large as the 95th percentile of the null distribution. The null distribution was calculated by 1, 000 permutations, as illustrated in Section 2. The type I error rate was estimated by calculating the power from data generated under the null hypothesis *H*_0_: *r*(*Z*) = *c*, which can be obtained by modifying the simulations where the random effect is set to be independent of *Z*. Theoretically, the type I error rate of the test should be equal to the significance level *α* = 0.05.

Table 1 shows the power calculated for eight different relationships with a varying sample sizes of 100, 200, and 500. Although both of the tested methods have low power in detecting modulators from a small sample size (*n* = 100), their power increases with the sample size. Note that GIMLET has higher power than MINDy in all relationships, except for the circle. Both GIMLET and MINDy have low chances of identifying modulators in the high-frequency sine relationship. GIMLET was shown to outperform MINDy, especially in the settings *M*_2_ and *M*_3_, because MINDy is not designed as a multivariate dependence measure for identifying modulators. Table 2 shows the type I error rates for the eight different relationships with varying sample sizes of 100, 200, and 500. The type I error rates are quite close to the chosen *α* level for all the tests, demonstrating that GIMLET shows better type I error rate control than MINDy, in this scenario.

**Table 1.** Statistical power of GIMLET and MINDy using synthetic data with different sample sizes (*n* = 100, 200, 500) for eight relationships (linear, quadratic, cubic, sine period 1/2, sine period 1/8, *x*^1/4^, circle, and step), under three different settings (*M*_1_, *M*_2_, and *M*_3_). The average of the *p*-values below the significant level *α* = 0.05 were calculated through 1,000 simulations.

| $n$ | Relationship | Simulation Model | | | | | |
| --- | --- | --- | --- | --- | --- | --- | --- |
| | | $M_1$ | | $M_2$ | | $M_3$ | |
|  |  | GIMLET | MINDy | GIMLET | MINDy | GIMLET | MINDy |
| 100 | Line | **0.895** | 0.576 | 0.621 | 0.090 | 0.652 | 0.141 |
|  | Quadratic | 0.536 | 0.307 | 0.219 | 0.062 | 0.663 | 0.140 |
|  | Cubic | 0.506 | 0.158 | 0.176 | 0.047 | 0.083 | 0.015 |
|  | Sine period 1/2 | 0.345 | 0.271 | 0.164 | 0.065 | 0.141 | 0.041 |
|  | Sine period 1/8 | 0.058 | 0.018 | 0.072 | 0.038 | 0.068 | 0.041 |
| | $x^{1/4}$ | 0.750 | 0.314 | 0.241 | 0.055 | 0.708 | 0.200 |
|  | Circle | 0.053 | 0.134 | 0.056 | 0.044 | 0.173 | 0.134 |
|  | Step | **0.880** | 0.554 | 0.545 | 0.089 | 0.364 | 0.041 |
| 200 | Line | **0.995** | **0.939** | **0.913** | 0.121 | **0.930** | 0.315 |
|  | Quadratic | **0.861** | 0.780 | 0.480 | 0.081 | **0.935** | 0.353 |
|  | Cubic | 0.767 | 0.520 | 0.334 | 0.039 | 0.235 | 0.025 |
|  | Sine period 1/2 | 0.679 | 0.463 | 0.336 | 0.061 | 0.297 | 0.040 |
|  | Sine period 1/8 | 0.078 | 0.013 | 0.105 | 0.027 | 0.123 | 0.019 |
| | $x^{1/4}$ | **0.939** | 0.693 | 0.474 | 0.043 | **0.949** | 0.405 |
|  | Circle | 0.128 | 0.342 | 0.078 | 0.019 | 0.420 | 0.250 |
|  | Step | **0.994** | 0.793 | **0.866** | 0.096 | 0.705 | 0.087 |
| 500 | Line | **1.000** | **1.000** | **1.000** | 0.208 | **1.000** | 0.761 |
|  | Quadratic | **0.995** | **1.000** | **0.885** | 0.090 | **1.000** | **0.822** |
|  | Cubic | **0.981** | **0.979** | 0.717 | 0.030 | 0.559 | 0.024 |
|  | Sine period 1/2 | **0.934** | **0.997** | 0.759 | 0.087 | 0.709 | 0.067 |
|  | Sine period 1/8 | 0.183 | 0.008 | 0.175 | 0.004 | 0.289 | 0.017 |
| | $x^{1/4}$ | **1.000** | **0.996** | **0.839** | 0.035 | **1.000** | **0.850** |
|  | Circle | 0.453 | **0.905** | 0.172 | 0.028 | **0.916** | 0.650 |
|  | Step | **1.000** | **0.998** | **0.997** | 0.143 | **0.965** | 0.202 |

**Table 2.** Type I error rate of GIMLET and MINDy, using synthetic data with different sample sizes (*n* = 100, 200, 500), for eight relationships (linear, quadratic, cubic, sine period 1/2, sine period 1/8, *x*^1/4^, circle, and step), under three different settings (*M*_1_, *M*_2_, and *M*_3_). The type I error rate of a test should be equal to the significance level *α* = 0.05.

| $n$ | Relationship | Simulation Model | | | | | |
| --- | --- | --- | --- | --- | --- | --- | --- |
| | | $M_1$ | | $M_2$ | | $M_3$ | |
|  |  | GIMLET | MINDy | GIMLET | MINDy | GIMLET | MINDy |
| 100 | Line | **0.047** | **0.042** | **0.045** | 0.071 | **0.057** | 0.061 |
|  | Quadratic | **0.041** | 0.029 | **0.052** | 0.076 | 0.064 | 0.063 |
|  | Cubic | **0.050** | 0.036 | **0.051** | 0.062 | 0.062 | **0.051** |
|  | Sine period 1/2 | 0.038 | 0.037 | **0.058** | 0.062 | **0.053** | **0.053** |
|  | Sine period 1/8 | **0.045** | 0.015 | **0.041** | **0.051** | **0.057** | **0.040** |
| | $x^{1/4}$ | **0.046** | 0.023 | **0.042** | 0.061 | **0.055** | **0.041** |
|  | Circle | 0.039 | 0.031 | **0.056** | **0.055** | **0.049** | **0.056** |
|  | Step | **0.053** | **0.049** | **0.048** | 0.077 | **0.054** | **0.056** |
| 200 | Line | **0.047** | 0.035 | **0.058** | 0.066 | **0.058** | 0.034 |
|  | Quadratic | 0.035 | 0.016 | **0.052** | **0.059** | 0.037 | **0.044** |
|  | Cubic | **0.045** | 0.021 | **0.048** | **0.046** | **0.043** | 0.027 |
|  | Sine period 1/2 | 0.062 | 0.021 | 0.034 | **0.047** | **0.043** | 0.030 |
|  | Sine period 1/8 | **0.049** | 0.009 | **0.045** | 0.030 | 0.039 | 0.026 |
| | $x^{1/4}$ | **0.049** | 0.030 | **0.059** | **0.056** | **0.059** | **0.041** |
|  | Circle | **0.059** | 0.017 | **0.048** | **0.056** | **0.046** | 0.035 |
|  | Step | **0.046** | 0.028 | 0.063 | **0.058** | **0.055** | 0.028 |
| 500 | Line | **0.047** | 0.021 | **0.041** | 0.030 | **0.040** | 0.012 |
|  | Quadratic | **0.051** | 0.017 | **0.053** | 0.023 | **0.046** | 0.012 |
|  | Cubic | **0.046** | 0.010 | **0.048** | 0.022 | **0.047** | 0.007 |
|  | Sine period 1/2 | **0.060** | 0.010 | **0.053** | 0.016 | 0.030 | 0.005 |
|  | Sine period 1/8 | **0.053** | 0.007 | **0.053** | 0.004 | **0.045** | 0.006 |
| | $x^{1/4}$ | **0.045** | 0.018 | **0.045** | 0.024 | **0.040** | 0.013 |
|  | Circle | **0.056** | 0.004 | **0.049** | 0.021 | **0.053** | 0.010 |
|  | Step | **0.046** | 0.012 | **0.044** | 0.023 | **0.047** | 0.012 |

## 4 Results with real data

We first sought to identify the genetic alterations that modulate the strength of the functional connection between HIF1A and the expression of its target genes in pan-kidney cohort in TCGA project [1]. The transcription factor HIF1A is a master transcriptional regulator of cellular and systemic homeostatic response to hypoxia. HIF1A activates the transcription of genes that are involved in crucial aspects of cancer biology, including angiogenesis, cell survival, glucose metabolism and invasion, and is implicated in the development of clear cell renal clear cell carcinoma (ccRCC). We examined mRNA expression profiles of 536 ccRCC and 357 non-ccRCC (papillary RCC and chromophobe RCC) patients, somatic mutation profiles of 436 ccRCC and 348 non-ccRCC patients, and copy number profiles of 528 ccRCC and 354 non-ccRCC patients, which can be downloaded from the Broad GDAC Firehose website [17]. We used 90 literature-validated target genes of HIF1A from the Ingenuity Knowledge Base [18] and calculated the factor scores for each patient by performing maximum-likelihood single factor analysis on the expression data matrix of these genes. In this example, we considered the factor score as the unobserved activity of HIF1A at the protein level and used it as *Y*. As candidates of *Z*, we first tested the somatic mutation of 85 genes, which were detected in more than 50 patients by genomic analyses of the pan-kidney cohort. We next considered the copy-number alterations of 41 chromosomal arms as candidates of *Z*. For this analysis, we expected to find an alteration of von Hippel-Lindau (VHL) tumor suppressor gene, which leads to overexpression of HIF1A and is a critical event in the pathogenesis of most ccRCC [19].

The modulator analysis of GIMLET yields five significantly associated gene mutations and genetic alterations modulating HIF1A activity with *q*-value<0.10 (Table 3). Indeed, GIMLET identified VHL as the most significantly associated gene mutation. Although PBRM1, identified as the second-most significantly associated gene mutation, is not reported to directly modulate HIF1A activity, this result remains significant because almost all PBRM1 mutant cases also have dysregulation of the hypoxia signaling pathway [**?**] and it is likely that PBRM1 and VHL cooperate in kidney carcinogenesis, which leads to the overexpression of hypoxia-inducible HIF1A. The analysis also yields three regions significantly modulating HIF1A activity with a *q*-value<0.10. Chromosome 3p deletions are observed in approximately 90% of ccRCC, which harbors VHL and tumor suppressor genes [21].

**Table 3.**
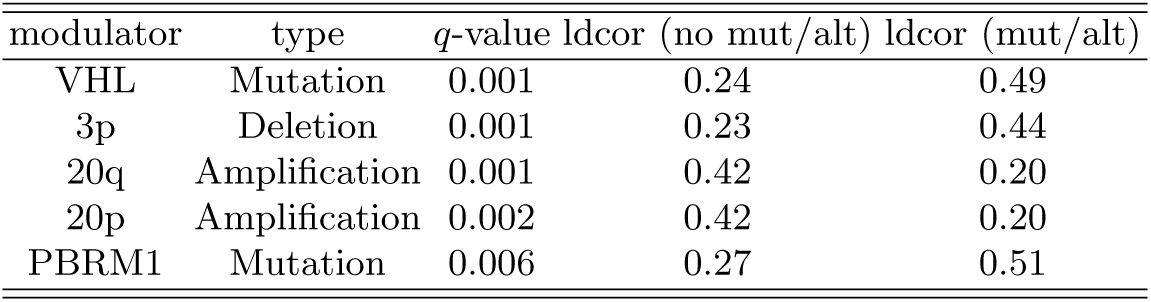
Five significantly associated gene mutations and genetic alterations modulating HIF1A activity.

We next examined drug-treated gene expression profiles from Broad Institute The Library of Integrated Cellular Signatures (LINCS) Center for Transcriptomics [22]. We sought to use these data to identify drugs that inhibit the strength of the functional connection between FOXM1 and CENPF which are master regulators of prostate cancer malignancy [23] and the expression of their target genes. A total of perturbational gene expression profiles of 22,268 probes for 6,684 experiments treated with 271 compounds after 24 h under different doses (0.04, 0.12, 0.37, 1.11, 3.33, and 10 *µ*m) in the two prostate cancer cell lines, PC3 and LNCaP, were downloaded from the LINCS L1000 dataset [22]. The expression values for each profile were normalized by robust *z*-scores relative to the control (plate population) and summarized using the median across replicates. If there are multiple probes that correspond to the same gene, the probe with the highest variance across all samples was selected as a single representative probe. Finally, the expression matrix data of 12,716 genes and 1,976 samples were used for further analysis. We used the expression of FOXM1 and CENPF as *X* and their unobserved activity as *Y* which was defined using maximum-likelihood single factor analysis on the expression data matrix for the 173 and 55 literature-validated targets of FOXM1 and CENPF, respectively, from the Ingenuity Knowledge Base [18]. The drug target genes for each compound under a given dose level were defined as differentially expressed genes, which were significantly lower in the drug-treated cell lines than in the vehicle-treated cell lines using a one-tailed t-test (*p*-value<0.001). As candidates of *Z*, the drug-perturbational activity for each sample under each of 1,850 different pertubagens was then estimated using the enrichment scores (maxmean statistics) of these drug target gene sets for gene set analysis [24]. We applied GIMLET to identify functional pertubagens modulating FOXM1 and CENPF activity.

The analysis yields 13 pertubagens that significantly inhibit the regulation of FOXM1 and CENPF with a *q*-value< 10^−7^ (Table 4). Indeed, these pertubagens support the inhibition of tumor progression in human prostate cancer by several resent studies. For example, Vorinostat known as suberanilohydroxamic acid is a member of a larger class of compounds that inhibit histone deacetylases (HDAC) [25]. A previous study has also shown that Vorinostat may inhibit tumor growth by both oral and parenteral administration in prostate cancer [26]. Withaferin A, a major bioactive component of the Indian herb Withania somnifera, induces cell death and inhibits tumor growth in human prostate cancer [27]. The activation of the PI3K-AKT-mTOR pathway is extremely common, if not universal, in castrate-resistant prostate cancer [28]. Certain PI3K and mTOR inhibitors are currently under investigation in clinical trials for CRPC including the dual inhibitor NVP-BEZ235 [29] and the mTOR inhibitor RAD001 or everolimus [30,31].

**Table 4.**
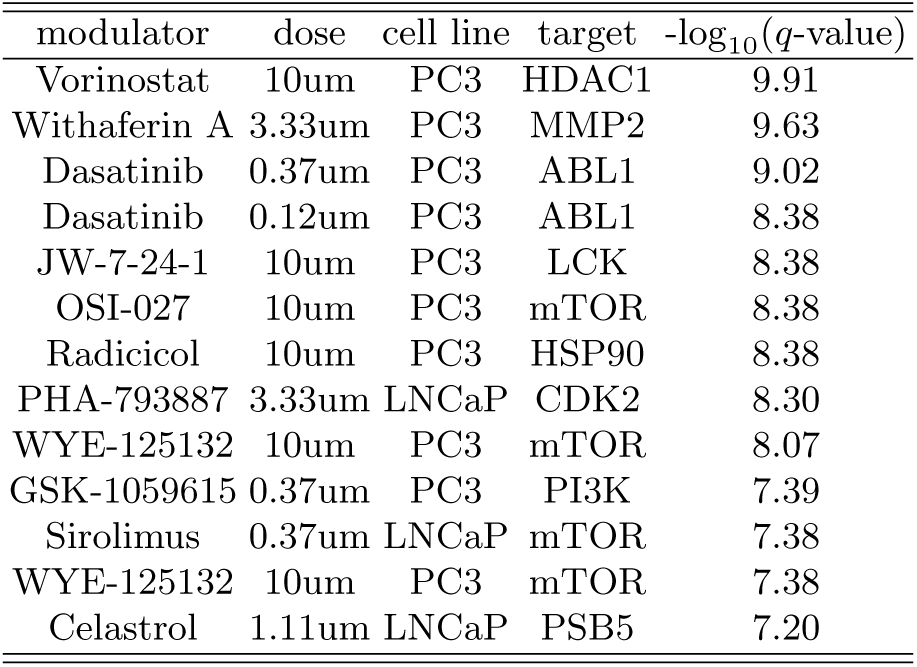
Thirteen significantly associated modulators (pertubagens) modulating FOXM1 and CENPF activity.

The analyses with two examples thus show that GIMLET can identify genetic alterations and functional pertubagens modulating the relationship between a given set of regulators and the expression of their target genes in particular cancer subtypes.

## 5 Discussion

The identification of modulators is a challenging problem for researchers who study gene regulation. The paradigm introduced by [4] and the state-of-the-art classical methods for identifying modulators are quite useful because they allow us to identify content-specific modulators of a TF activity using gene expression data. However, these methods are restricted to the capturing of a particular type of dependency between univariate random variables, and it can be difficult to describe more complex multivariate dependency structures, when TFs and modulators are functionally related. We have developed a more general class of the identification of modulators, in the framework of energy statistics and a specific implementation, called GIMLET. An appealing property of the proposed method is that it can easily measure all types of dependencies, including nonmonotonic and nonlinear relationships, between random vectors in an arbitrary dimension. Our simulation results demonstrate that GIMLET outperforms MINDy in terms of its statistical power and type I error rate. An analysis with a real example thus showed that GIMLET can identify genetic alterations and functional pertubagens modulating TF activities. We believe that the presented method may be useful for a range of biological applications, and it could represent a breakthrough in gene regulation research.

## Acknowledgement

This work was supported by JSPS Grant-in-Aid for Challenging Exploratory Research (15K12139), JSPS Grant-in-Aid for Young Scientists A (15H05325), and JSPS Grant-in-Aid for Scientific Research on Innovative Areas (15H05912 and 18H04798). It was also supported in part by Ministry of Education, Culture, Sports, Science and Technology (MEXT) of Japan as a social and scientific priority issue (Integrated computational life science to support personalized and preventive medicine; hp170227, hp180198) to be tackled by using post-K computer. The super-computing resources were provided by Human Genome Center, University of Tokyo.

